# Evolution of sex hormone binding globulins reveals early gene duplication at the root of vertebrates

**DOI:** 10.1101/2020.07.02.184432

**Authors:** Yann Guiguen, Jeremy Pasquier, Alexis Fostier, Julien Bobe

**Affiliations:** INRAE, LPGP, 35000 Rennes, France

**Keywords:** evolution, Sex hormone-binding globulin, teleosts, vertebrates, gene expression

## Abstract

Sex hormone-binding globulin (Shbg) is an important vertebrate blood carrier protein synthetized in the liver and involved in the transport and local regulation of sex steroids in target tissues. A novel *shbg gene (shbgb)* with a predominant ovarian expression was recently characterized. Being initially found only in salmonids, this *shbgb* was originally thought to result from the Salmonid-specific whole genome duplication. Using updated transcriptomic and genomic resources we identified *Shbgb* orthologs in non-salmonid teleosts (European eel, arowana), holosteans (spotted gar, bowfin), polypteriformes (reedfish), agnatha (sea lamprey) and in amphibians, and found that the classical *Shbg* gene (*Shbga*) displays a predominant hepatic expression whereas *Shbgb* has a predominant gonadal expression. Together, these results indicate that these two *Shgb* genes most likely originate from a whole genome duplication event at the root of vertebrate evolution, followed by numerous and independent losses and by tissue expression specialization of *Shbga* and *Shbgb* paralogs.

**Highlights:**

- Phylogeny, synteny and expression analyses shed new light on Shbg evolution in vertebrates.
- Shbg diversity originates from a duplication event at the root of vertebrate evolution.
- This duplication was followed by many independent losses of *Shbg* paralogs in vertebrates.
- *Shbg* paralogs have acquired different tissue expression patterns.

## 1 Introduction

Sex hormone-binding globulin (Shbg) is mainly known as a blood-protein carrier involved in the transport of sex steroids in the plasma and in the regulation of their bioavailability to target organs. Shbg proteins present sequence similarities with the other LamG domain-containing proteins growth arrest-specific 6 (Gas6) and protein S alpha (Pros1) (Joseph, 1997; Joseph and Baker, 1992). By transporting and regulating androgens and estrogens access to the gonads, Shbg plays important roles in vertebrates reproduction (Hammond, 2011). The Shbg protein was originally identified in the beta-globulin fraction of the human serum (Rosner et al., 1969) and has previously been known as Androgen-binding protein (Abp). *Shbg* genes and Shbg proteins have been characterized in a variety of tetrapod species with the notable exception of birds (Westphal, 1986; Wingfield et al., 1984). In aquatic vertebrates, Shbg was originally found in the plasma of an elasmobranch, the skate (*Raja radiata*) (Freeman and Idler, 1969), and of a teleost, the rainbow trout (*Oncorhynchus mykiss*) (Fostier and Breton, 1975). Since that moment on, Shbg have been subsequently identified and studied in many fish species (see for review (Bobe et al., 2010)). Two different *shbg* genes i.e., s*hbga* and *shbgb*, have been characterized in teleosts with *shbga* being the ortholog of the mammalian *Shbg*, which has been conserved from chondrichthyes to tetrapods and *shbgb* that has only been reported up to now in salmonids. In contrast to *shbga* that is mainly expressed in the liver (Bobe et al., 2008), the *shbgb* transcript was mainly found in the ovary, suggesting a local mediation of the sex steroids effects by the Shbgb protein (Bobe et al., 2008). These two salmonid Shbg proteins share very low identity percentages, with for instance 26% identity between Shbga and Shgbb at the amino acid level in the rainbow trout (Bobe et al., 2010, 2008). In comparison to other vertebrates and teleost fishes, the salmonid ancestor has experienced an additional (4R) whole genome duplication known as the salmonid specific whole genome duplication, or SaGD (Berthelot et al., 2014). For this reason, and because no *shbgb* gene had ever been reported in a non-salmonid species, it was first hypothesized that *shbga* and *shbgb* were ohnologous genes resulting from the SaGD (Bobe et al., 2008). This hypothesis was subsequently challenged in another study and the possibility of an ancient duplication followed by a lineage-specific retention in salmonids was suggested (Miguel-Queralt et al., 2009).

The evolutionary history of *Shbg* genes in vertebrates thus remained unclear and deserved further investigations. Using the increasing amount of genomic and transcriptomic data available for many vertebrate species we revisited the evolutionary history of *Shbg* genes. The transcriptomes of 24 actinopterygian species (including 22 teleosts) and vertebrate genomes were included in the analysis, which led to the identification of previously non-characterized *Shbgb* genes in several non-salmonid vertebrate lineages. Using phylogenomic analyses, we identified several *Shbgb* orthologs in a variety of non-salmonid vertebrate species, including teleosts, non-teleost actinopterygians, amphibians and one agnatha. Combined with synteny reconstruction analysis, we demonstrated that *Shbg* diversity results from a duplication event much older than the SaGD. To gain new information on the functional evolution of *shbg* genes, we also used quantitative PCR and next generation sequencing approaches, to characterize the expression profiles of *shbga* and *shbgb* transcripts for several actinopterygian species. This showed that the paralogous *shbg* genes have acquired different expression profiles with *shbgb* having a predominant gonadal expression contrasting with a predominant liver expression of *shbga*.

## 2 Material and Methods

### 2.1 Genomic and transcriptomic databases

The genomes of the following species, human, *Homo sapiens*; tropical clawed frog, *Xenopus tropicalis*; coelacanth, *Latimeria chalumnae; spotted gar, Lepisosteus oculatus and zebrafish, Danio rerio* were explored using the Ensembl genome browser (http://www.ensembl.org/index.html). The rainbow trout (*Oncorhynchus mykiss*) genomic database was searched using the Genoscope trout genome browser (http://www.genoscope.cns.fr/trout/). The European eel (*Anguilla Anguilla*) genomic database was investigated using the European eel assembly available at ZF-Genomics (http://www.zfgenomics.org/sub/eel). Transcriptomes of holostean and teleostean species were investigated using the PhyloFish project resource (Pasquier et al., 2016) available at http://phylofish.sigenae.org. The protein sequences of Human SHBG, *Xenopus* Shbg, zebrafish Shbg, and rainbow trout Shbga and Shbgb were used as queries to identify homologs of Shbga and Shbgb in the different genomic and transcriptomic databases investigated. A similar methodology was used for Gas6 and Pros1 proteins that were relevant to study due to their phylogenetical proximity and structural similarity.

### 2.2 Phylogenetic and synteny analyses

Amino-acid sequences of 126 predicted Shbg (a, b), Gas6 and Pros1 proteins were first aligned using ClustalW (Thompson et al., 1994), then alignments were manually adjusted, to improve the quality of the multiple sequence alignments. The JTT (Jones, Taylor and Thornton) protein substitution matrix of the resulting alignment was determined using ProtTest software (Darriba et al., 2011). Phylogenetic analysis of the proteins presenting LamG domains (*i*.*e*. Shbga, Shbgb, Gas6 and Pros1) was performed using the neighbour joining (NJM) method (MEGA 5.1 software), with 1000 bootstrap replicates (Tamura et al., 2011). Trees were edited online with iTOL (Letunic and Bork, 2016) and exported as Scalable Vector Graphics.

Synteny maps of the conserved genomic regions in human, *Xenopus*, coelacanth, spotted gar and zebrafish were constructed based on information available within the Genomicus (Muffato et al., 2010) v75.01 website (http://www.genomicus.biologie.ens.fr/genomicus-75.01/cgi-bin/search.pl). Synteny map of the conserved genomic regions in the rainbow trout was performed using the Rainbow Trout Genomicus Server (http://www.genomicus.biologie.ens.fr/genomicus-trout-01.01/cgi-bin/search.pl). The synteny analyses of European eel conserved genomic regions were obtained performing TBLASTN searches in the corresponding genomic database. For each studied gene, the protein sequences of human and zebrafish were used as queries.

Multiple alignments plots of *shbgb* genes in salmonids were processed online (http://genome.lbl.gov/vista/) with mVISTA (Dubchak and Ryaboy, 2006; Poliakov et al., 2014) using genomic *shbgb* sequences of rainbow trout, *Oncorhynchus mykiss*, Atlantic salmon, *Salmon salar*, and Coho salmon, *Oncorhynchus kisutch*. Putative *shbgb* pseudogenes were retrieved by TBLASTN searches on whole genome sequences using as query the protein sequence of rainbow trout Shbgb.

### 2.3 RNA-seq *shbga* and *shbgb* tissue expression in holosteans and teleosts

RNA-seq and *de novo* assembly were performed for all studied species as previously described (Berthelot et al., 2014; Braasch et al., 2016; Pasquier et al., 2016). In order to study the expression patterns and levels of *shbg* transcripts for each actinopterygian species with two *shbg* genes, we mapped RNA-seq reads on the corresponding *shbg* coding sequence (CDS) using BWA-Bowtie (Langmead and Salzberg, 2012) with stringent mapping parameters (maximum number of allowed mismatches –aln 2). Mapped reads were counted using SAMtools (Li et al., 2009) idxstat command, with a minimum alignment quality value (– q 30) to discard ambiguous mapping reads. For each species, the numbers of mapped reads were then normalized for each *shbg* gene across the eleven tissues using the reads per kilo base per million mapped reads (RPKM) normalization. All RNA-seq data are available here: (http://phylofish.sigenae.org/index.html)

### 2.4 Quantitative PCR analysis (QPCR)

QPCR was performed using the RNA collections of the PhyloFish RNA-seq project as previously described (Braasch et al., 2016; Pasquier et al., 2016). Briefly, tissues were sampled from the same female individual and testis from a male individual, when possible. In some species and depending on the tissues, RNA samples from different individuals were pooled to obtain sufficient amounts of RNA. Total RNA was extracted using Tri-Reagent (Molecular Research Center, Cincinnati, OH, USA) according to the manufacturer’s instructions. Reverse transcription (RT) was performed using 1 μg of RNA for each sample with M-MLV reverse transcriptase and random hexamers (Promega, Madison, WI, USA). Briefly, RNA and dNTPs were denatured for 6 min at 70°C, chilled on ice for 5 min before the RT master mix was added. RT was performed at 37°C for 1 h and 15 min followed by a 15-min incubation step at 70°C. Control reactions were run without reverse transcriptase and used as negative control in the real-time PCR study. Quantitative RT-PCR (QPCR) experiments were performed sing an Applied Biosystems StepOne Plus. RT products, including control reactions, were diluted to 1/25, and 4 μl were used for each PCR. All QPCR were performed in triplicates. QPCR was performed using a real-time PCR kit provided with a Fast-SYBR Green fluorophore (Applied Biosystems) with 200 nM of each primer in order to keep PCR efficiency between 80% and 100% for all target shbg genes. The relative abundance of target cDNA within a sample set was calculated from serially diluted cDNA pool (standard curve) using Applied Biosystem StepOne V.2.0 software. After amplification, a fusion curve was obtained to validate the amplification of a single PCR product. The fusion curves obtained showed that each primer pair used was specific of a single *shbg* transcript. The negative control reactions were used to estimate background level. Genes were considered significantly expressed when measured level was significantly above background at p□<□0.05 and within the range of the standard curve. For each studied tissue, cDNA originating from three individual fish were pooled and subsequently used for real-time PCR. Before further analysis, real-time PCR data were collected using the same detection threshold for all studied genes. Data were subsequently normalized using the ΔΔCt method to 18S transcript abundance in samples diluted to 1:2,000.

### 2.5 Clustering analysis

Expression profiles originating from either QPCR and RNA-seq were represented using supervised clustering methods (Eisen et al., 1998). Hierarchical clustering was processed using centroïd linkage clustering, that uses the average value of all points in a cluster as a reference to calculate distance of other points, with Pearson’s uncentered correlation as similarity metric on data that were normalized and median-centered using the Cluster program (Eisen et al., 1998). Results (colorized matrix) of hierarchical clustering analyses were visualized using the Java TreeView program (Saldanha, 2004).

## 3 Results

### 3.1 Shbg gene evolution in vertebrates

In order to decipher phylogenetic relationships among Shbg sequences, a phylogenetic reconstruction of the evolution of Shbg was made based using the alignment of 126 vertebrate LamG domain-containing proteins. This phylogeny includes Shbg proteins, growth arrest specific 6 proteins (Gas6) and Vitamin K-dependent protein S (Pros1) and the tree was rooted using as outgroup the zebrafish Laminin subunit alpha 4 (lama4). This analysis (Fig. 1) shows that these vertebrate LamG domains proteins cluster into two major clades containing Shbg proteins on one side and the Gas6 and Pros1 proteins on the other side. These Shbg and Gas6/Pros1 clades are both significantly supported with high bootstrap values (*i*.*e*. 75% and 100%, respectively). The Shbg clade contains two sub-clades both supported by significant bootstrap values. The Shbga cluster (100% bootstrap support), in red, contains all classical vertebrate Shbg proteins from chondrichthyes to teleosts with the notable exception of birds in which no Shbg proteins have been detected (see also Fig.S2). The Shbgb cluster (93% bootstrap support), in blue, contains not only the salmonid Shbgb proteins, but also other vertebrate sequences outside the salmonid family including sequences from teleosts (European eel and silver arowana), non-teleost bony fishes (reedfish, spotted gar and bowfin), some amphibians and an agnatha, i.e., the sea lamprey (Fig.1 and Fig.S1). The tree topology indicates that Shbgb proteins are not specific to the salmonids lineage and thus suggests a much more ancient origin of *Shbgb* genes in vertebrates than previously hypothesized (Bobe et al., 2008).

**Figure 1:**
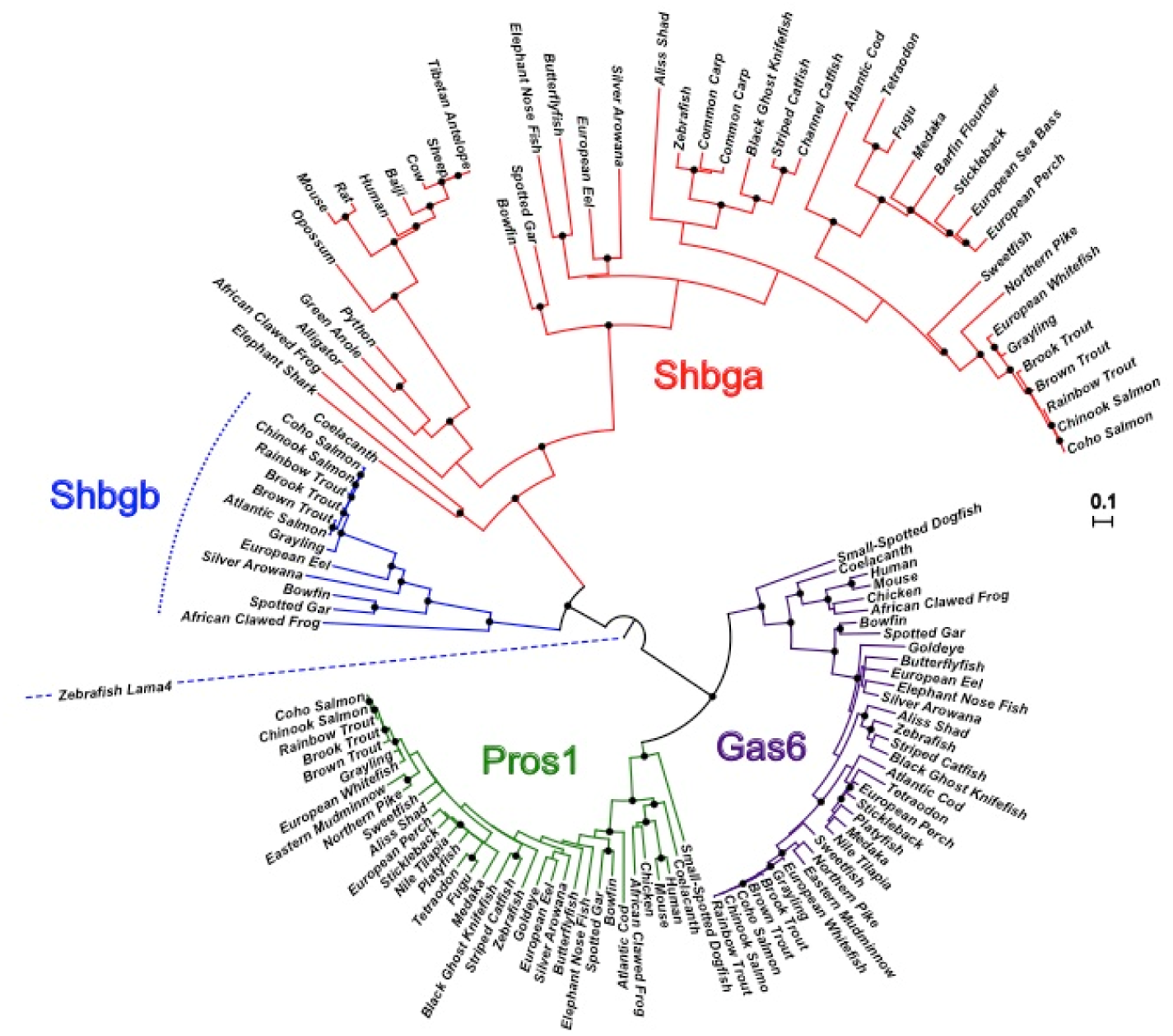
Phylogenetic reconstruction of the evolution of LamG domains proteins (Shbg, Gas6 and Pros1) in vertebrates. Circular NJM phylogenetic tree of LamG domains proteins including Shbga in red, Shbgb in blue, Gas6 in purple and Pros1 in green. The tree is rooted using zebrafish Laminin Subunit Alpha 4 (lama4) and bootstrap values over 0.75 are shown with a black dot on each significant node.

To strengthen this phylogeny-based analysis of Shbg protein evolutionary history we carried out a synteny analysis in order to better support this hypothesis of an ancient origin *Shbga* and *Shbgb* genes. The synteny analysis first revealed that these two *Shbg* genes are located on two different syntenic chromosome regions in vertebrates (see Fig. 2A for the *Shbga* locus and Fig. 2B for the *Shbgb* locus). In contrast, *Gas6* and *Pros1* genes are located on the same syntenic chromosome region in most studied species, with the exception of primates (Fig. 2C). In vertebrates, *Shbga, Shbgb*, and *Gas6*/*Pros1* are also located in regions containing other syntenic gene families. These neighboring genes are spread over four syntenic regions (Fig. 2A-2D) like for instance for the *Atp*-related genes (*Atp1b2, Atp1b3, Atp4b*) that are found in all four syntenic chromosome regions. In addition, *Zbtb*-related genes (*Ztb4, Zbt38, Zbt33*) and *Lamp*-related genes (*Lamp1, Lamp2, Lamp3*) are present in three of the four different syntenic chromosome regions depending of the gene family. Altogether, these results strongly suggest that the diversity of the gene family present on these four syntenic chromosome regions probably results from early whole genome duplication events (VG1 and/or VG2) that occurred at the root of vertebrate evolution with a subsequent complex pattern of gene retention and gene losses.

**Figure 2:**
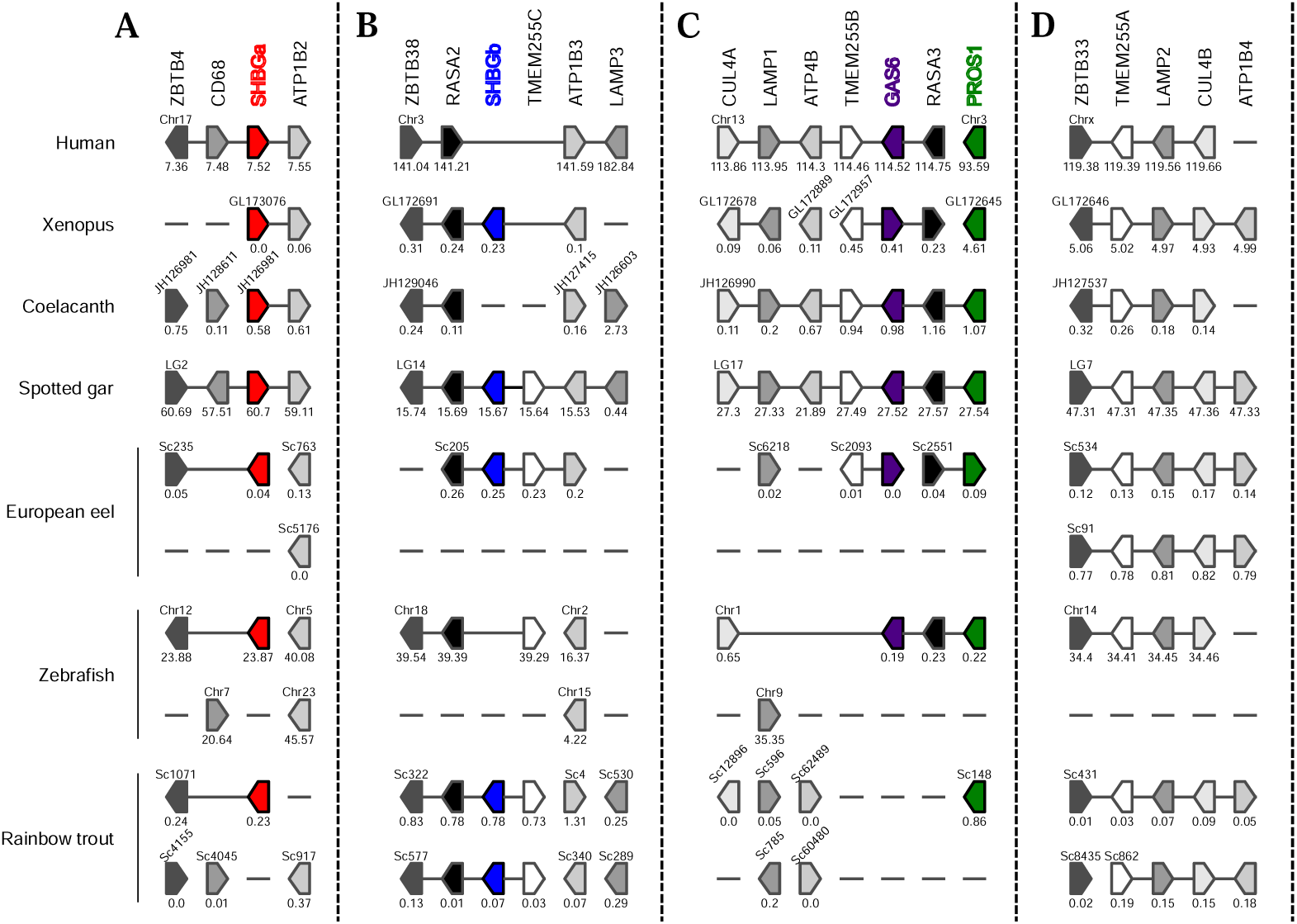
Synteny maps of conserved genomic regions around LamG domains proteins (i.e. Shbga, Shbgb, Gas6 and Pros1) in human, Xenopus, coelacanth, spotted gar, Euroean eel, zebrafish and rainbow trout. Synteny maps are given for genomic regions around *Shbga* (**A**), *Shbgb* (**B**), *Gas6* and *Pros1* locus (**C**) and for a fourth region containing homologs of neighbouring *Shbga, Shbgb, Gas6* and *Pros1* genes (**D**). Genes are represented by blocks with an arrowed side indicating the gene orientation on chromosomes, linkage group or scaffolds. Gene location on chromosomes (Chr for Human and zebrafish), Linkage group (LG for spotted gar) and scaffolds (Ensembl reference or scaffold number) is given in Mb below each gene block. Genes belonging to the same Chr, LG or scaffolds are linked by a solid line.

Using the recently released salmonid genome resources, we also re-investigated the presence of additional copies of *shbg* genes in salmonids and confirmed that *shbga* and *shbgb* were both retained as single copies (Fig. 1) in rainbow trout, Atlantic salmon and coho salmon suggesting that no functional duplicated copies were retained after the salmonid whole genome duplication (SaGD). However, we also found an additional *shbgb* gene conserved in these three salmonid species (Fig.3A), but with many stop codons in its deduced open reading frame (see example for rainbow trout in Fig.3B), suggesting that this gene *(ψ shbgb)* was subsequently pseudogenized after the SaGD.

**Figure 3:**
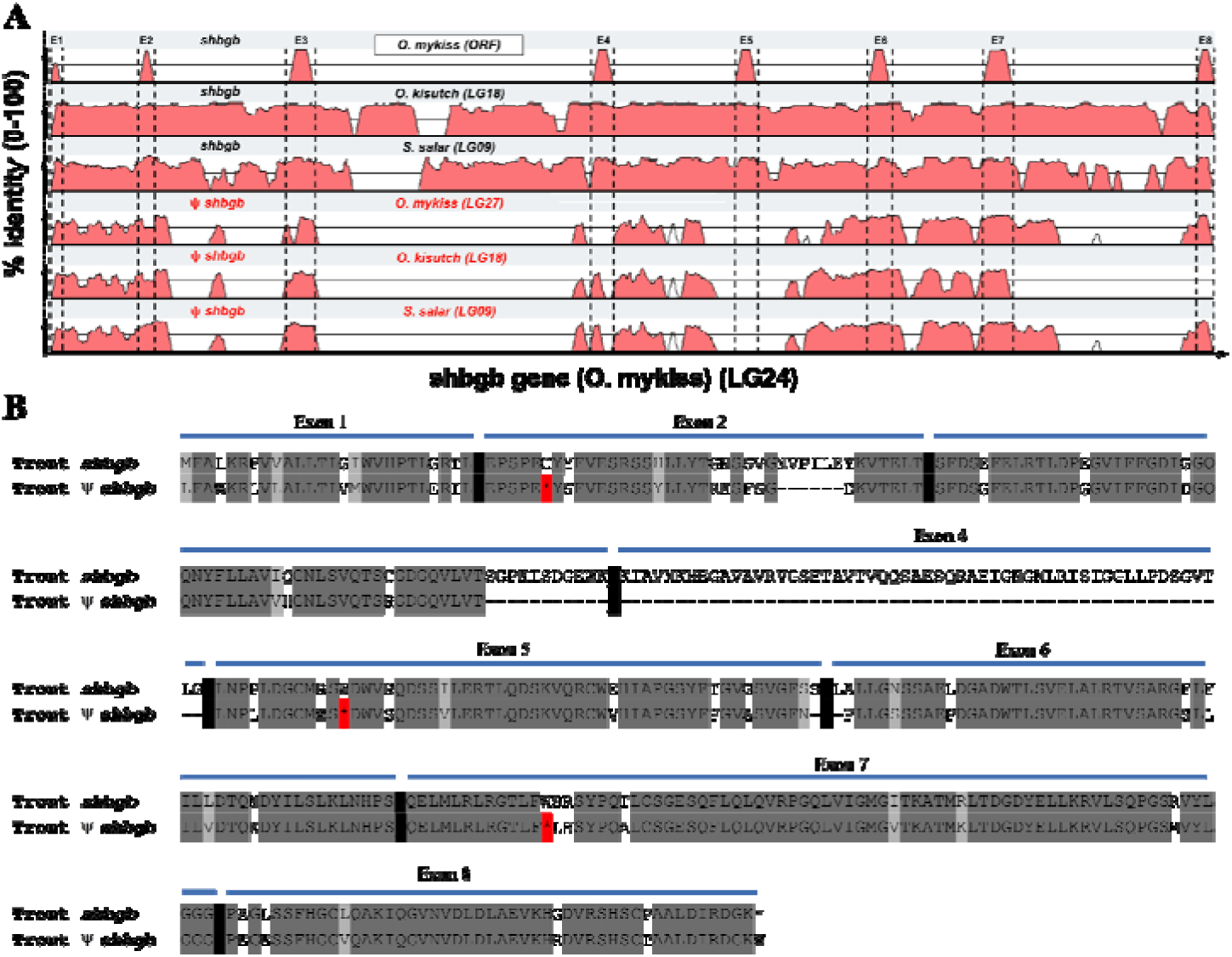
Multiple alignments plots of *shbgb* genes in salmonids. (**A**). Percentage of sequence identity of *shbgb* genes in Atlantic salmon, *Salmon salar*, and Coho salmon, *Oncorhynchus kisutch* compared to the rainbow trout, *Oncorhynchus mykiss, shbgb* gene on linkage group (LG) 24. In addition to the functional *shbgb* genes that were found in all salmonid species investigated, all these species have an additional *shbgb* homolog containing multiple stop codons and thus considered as a pseudogene *(ψ shbgb)*. (**B**). Rainbow trout Shbgb and ψ Shbgb protein alignment showing that the ψ Shbgb contains multiple stop codons (red asterisks) and a large deletion in exon 4 of the Shbgb protein.

### 3.2 Expression patterns of shbga and shbgb

In all investigated actinopterygians, *shbga* was found to be mainly expressed in the liver, supporting a conserved role for this blood-secreted Shbg (Fig.4). However, in the silver arowana a low *shbga* expression in the gonads is also detected in addition to the predominant liver expression (Fig.4A and Fig.4B). In contrast to *shbga*, expression of *shbgb* is not predominant in the liver in all investigated actinopterygians (Fig.4A and Fig.4B). In contrast, *shbgb* expression is predominantly detected in the gonads (ovary and/or testis) with the notable exception of the European eel.

**Figure 4:**
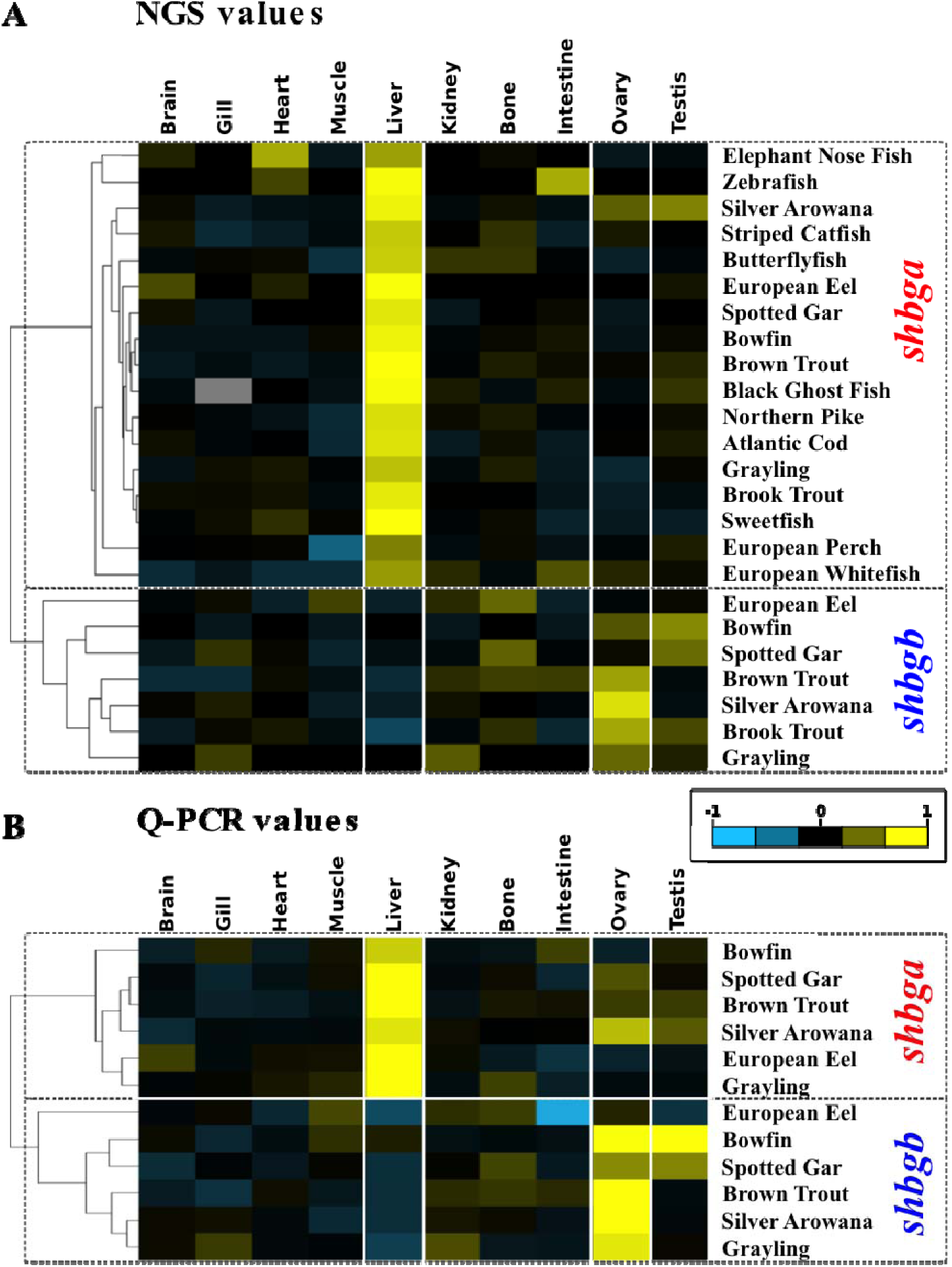
Tissue expression profiles of *shhbga and shbgb* genes. (**A**). Heatmap (colorized matrix) of the hierarchical clustering of tissue RNA-Seq expression profiles (NGS values) of *shbg* genes in different Holostean (spotted gar and bowfin) and teleost species. A predominant expression of *shbga* is found in the liver contrasting with the predominant expression of *shbgb* in gonads (**B**). Heatmap (colorized matrix) of the hierarchical clustering of tissue expression profiles of *shbg* genes analyzed by quantitative PCR (QPCR values) in different Holostean (spotted gar and bowfin) and teleost species. Colorized matrixes highlight the high expressing tissues in yellow, the low expressing tissues in blue and the median expression in black (see color scale).

**Figure 5:**
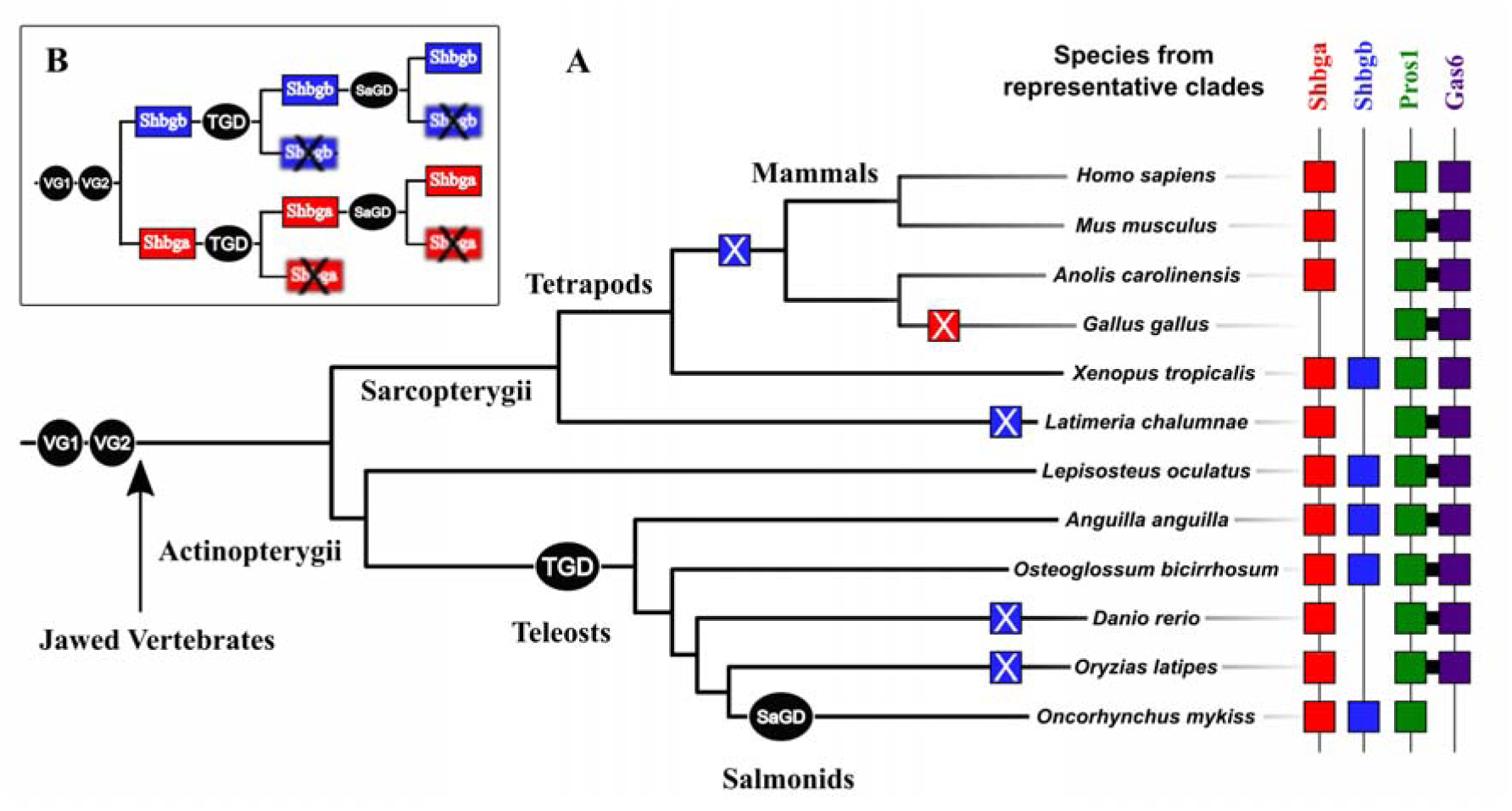
Evolution of *Shbg*/*shbg, Gas6*/*gas6*, and *Pros1*/*pros1* genes through successive whole genome duplications and independent gene losses. (**A**). Schematic representation of the evolution of LamG domains proteins including, Shbga (red squares), Shbgb (blue squares), Gas6 (purple squares) and Pros1 (green squares) in some jawed vertebrates’ representative species. Whole genome duplications (black circles; VG1 and VG2: vertebrate genome duplications 1 and 2, TGD: teleost genome duplication, SaGD: salmonid genome duplication) are indicated at each duplication nodes. Gene losses are represented by square boxes with a cross inside. (**B**). Simplified representation of the evolution of Shbg proteins after whole genome duplications showing the systematic losses of one Shbga and Shbgb paralog after each duplication.

## 4 Discussion

In this study, we aimed at investigating the diversity of the Shbg family in vertebrates and the evolutionary history of *Shbg* genes. To date, despite the recent discovery of a second *shbg* gene (i.e. *shbgb*) in salmonids, the origin and diversity of *Shbg* genes in vertebrates has remained controversial. Because salmonids experienced an additional whole genome duplication (SaGD) approximately 100 Mya (Berthelot et al., 2014; Macqueen and Johnston, 2014) compared to other teleost fish and as *shbgb* genes were initially found in salmonids and, never reported at that time in any non-salmonid species, *shbga* and *shbgb* have been first hypothesized to be the result of the SaGD (Bobe et al., 2010, 2008). However pairwise comparison of Shbga and Shbgb reveals a surprisingly low sequence identity (around 25% at the amino acid level) that was initially interpreted as Shbga and Shbgb being highly divergent SaGD paralog. However, this paralogy relationship was not supported by the phylogeny reconstruction (Bobe et al., 2008) and this discrepancy was thus explained as being the result of a long-branch attraction artifact resulting from the dramatic divergence of these two sequences (Bobe et al., 2010, 2008). Based on the cloning of another salmonid *shbgb* gene in coho salmon, *Oncorhynchus kisutch* and the low sequence identity between salmonids Shbga and Shbgb, other authors (Miguel-Queralt et al., 2009), hypothesized that the *shbgb* gene could stemmed from a much more ancient duplication than the SaGD. In order to decipher the evolutionary history of *Shbg* genes we re-analyzed the phylogenetic relationships of *Shbg* genes, their local synteny context, and the evolution of the phylogenetically and structurally closely related genes i.e., *Gas6* and *Pros1* that also contain LamG domains and are often identified as potential members of the same family. The identification of new *Shbgb* genes in vertebrates, the *Shbg* phylogenetic tree topology and their local synteny relationships strongly suggest that *Shbga* and *Shbgb* genes result from a whole genome duplication event that occurred very early at the root on the vertebrate lineage. The presence of a single *Shbga* gene and a single *Shbgb* gene in amphibians, holosteans, polypteriformes, agnatha and some teleost fishes, suggests that this *Shbg* duplication stems at least from the second round of vertebrate genome duplication (VG2). It is however also possible that *Shbga* and *Shbgb* originate the first round of vertebrate genome duplication (VG1) followed by the loss of one duplicate of each gene before VG2. Following this early duplication, these two *Shgb* paralogs have evolved through many different phylum-specific gene retentions and/or gene losses. Among them the case of birds is interesting as they not only lost their *Shbgb* gene like reported here for many other tetrapods, but also their *Shbga* gene that is found to be conserved in all other vertebrates. This complete absence of *Shbga* in birds has been already reported and it was hypothesized that this specific steroid hormone-binding transport would then be performed by a corticosteroid-binding globulin (Wingfield et al., 1984). Similarly, no Shbg homologs were detected by homology searches in Chondrichthyes (data not shown) but their complete absence in this clade requires further in-depth analysis and additional genome information as Shbg-like sex steroid binding capacities exist in the serum of the Thorny skate (Freeman and Idler, 1969). In tetrapods, Shbgb was only found in Amphibians along with Shbga. Shbgb was also found in holosteans (spotted gar and bowfin), polypteriformes (reedfish), agnatha (sea lamprey) and in a few teleost fish orders i.e., in Elopomorphs (European eel), Osteoglossiforms (silver arowana) and Salmoniforms even though the protein is frequently misannotated in GenBank (Fig.S1). Interestingly we did not find any retention of additional whole genome [SaGD and the teleost specific duplication (TGD)] paralogs for both *shbga* and *shbgb* gene with the exception of a pseudogenized SaGD *shbgb* paralog (*ψ shbgb*). This indicates that these extra whole genome duplications did no impact the repertoire of *shbg* genes with a maximum of one *shbga* and one *shbgb* functional copies in all investigated teleost clades. This systematic and independent losses of additional *shbga* and *shbgb* duplicated paralogs in teleosts may reflect an evolutionary constraint of maintaining a correct gene and protein dosage as it has been suggested in other organisms (Conant et al., 2014; Gout and Lynch, 2015).

In consistency with existing data in mammals, our expression data showed that *shbga* is predominantly expressed in the liver in the different teleost species studied here. This confirms what has previously been reported in various teleost species including zebrafish (*Danio rerio*) (Miguel-Queralt et al., 2004), rainbow trout (*Oncorhynchus mykiss*) (Bobe et al., 2008), Coho salmon (*Oncorhynchus kisutch*) (Miguel-Queralt et al., 2009), pejerrey (*Odontesthes bonariensis*) (González et al., 2017) and sea bass (*Dicentrarchus labrax*) (Miguel-Queralt et al., 2007). In addition, this strong hepatic expression is also observed in spotted gar (*Lepisosteus oculatus*) and bowfin (*Amia calva*) as shown by both RNA-seq and QPCR data.

In contrast to *shbga*, data on the tissue distribution of *shbgb* remain scarce. The ovarian predominant expression of *shbgb* was originally reported in rainbow trout, in which the transcript could also be detected at lower levels in muscle and stomach (Bobe et al., 2008). Semi quantitative data in Coho salmon confirmed the expression of *shbgb* in the ovary and stomach and revealed its presence in gills (Miguel-Queralt et al., 2009). Here we show that *shbgb* is also predominant expressed in the ovary in brown trout, silver arowana and grayling. We also report a strong testicular expression of *shbgb* in the two holostean species, spotted gar and bowfin, that appears to be lost in teleosts. In addition, the *shbgb* gene does not exhibit any gonad predominant expression in European eel. Together, our data show that *shbga* and *shbgb* have a very specific expression patterns with a predominant expression in liver and gonads, respectively. This pattern appears to be conserved during evolution without any significant change following whole genome duplications events (TGD and SaGD), with the exception of European eel in which the gonad predominant expression of *shbgb* appears to be lost. Finally, the strong testicular expression of *shbgb* revealed in bowfin and spotted gar is not found in any teleost species suggesting a specific role of Shbgb in testicular physiology in holostean species.

The multiple independent losses of *Shbgb* across vertebrates, while *Shbga, Gas6* and *Pros1* have been conserved in almost all vertebrates, could reflect different adapative and reproductive strategies as Shbg have been shown to be important carrier proteins for the blood transport of sex steroids and for their delivery to target reproductive tissues (Hammond, 2011). However, despite this discrepancy among species, the distinct roles of Shgba in hormone transport in the blood and of Shbgb in local hormone action in reproductive organs as well as the associated expression in liver and gonads, respectively, appears to be evolutionary conserved in species that have retained both genes despite a few intriguing species-specific exceptions.

## 5 Acknowledgement

This work was supported by the French national research Agency (ANR-10-GENM-017– PhyloFish).

## Supplementary Figures

**Supplementary Figure 1:**
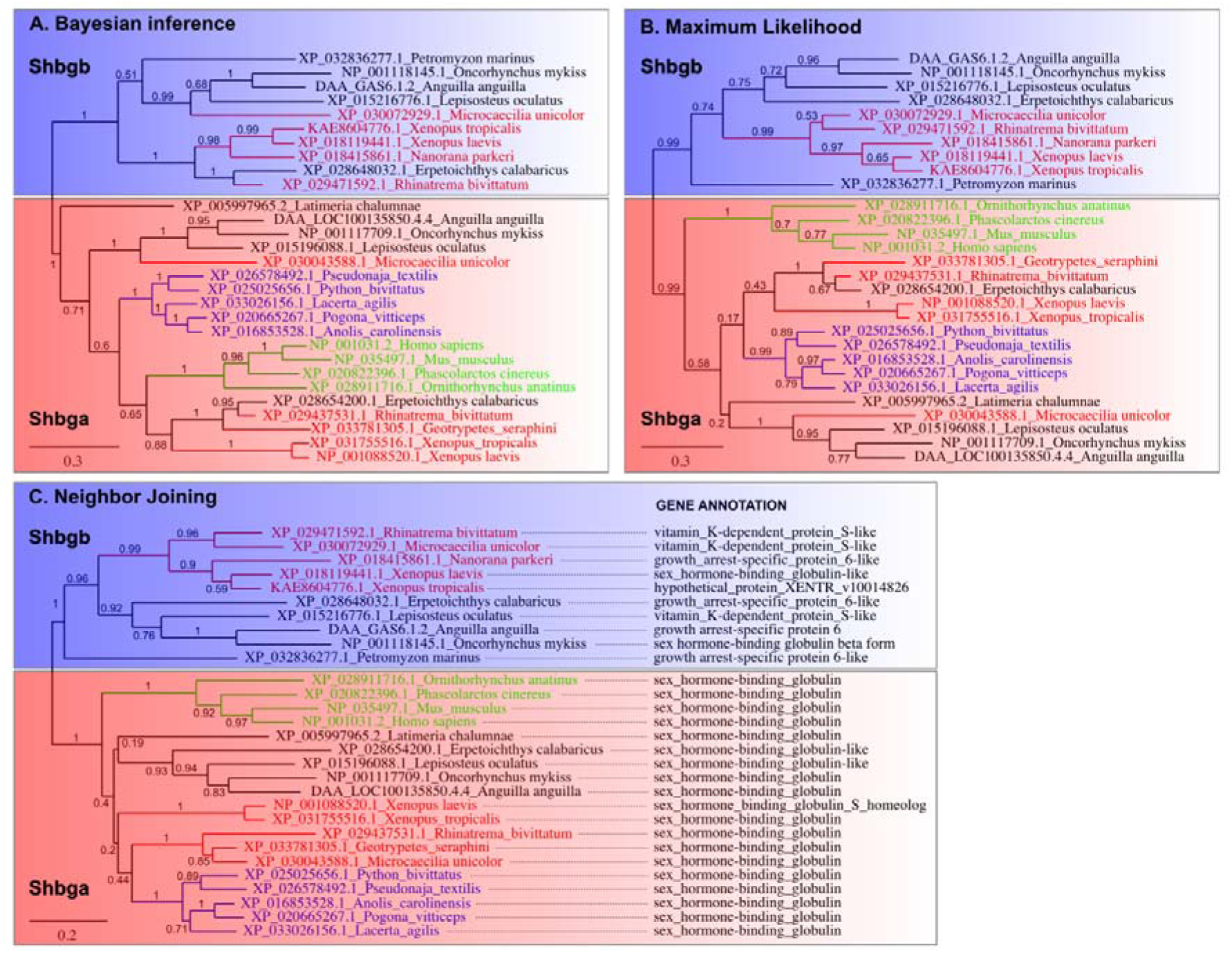
Phylogenetic trees of some vertebrate Shbga and Shbgb proteins. Protein sequences were searched in the ref-seq GenBank database using blastp (protein-protein BLAST) and in the the PhyloFish website (http://phylofish.sigenae.org/) using Shbga and Shbgb proteins from spotted gar (*Lepisosteus oculatus*, XP_015196088.1 and XP_015216776.1) and Western claw frog (*Xenopus tropicalis*, XP_0311755516.1 and KAE8604776.1) as baits. Phylogenetic analyses were performed on the Phylogeny.fr website (http://www.phylogeny.fr). Sequences were aligned with MUSCLE (v3.8.31) and cleaned with Gblocks (v0.91b) with default settings. Phylogenetic trees were reconstructed using the bayesian inference (**A**.) method implemented in the MrBayes program (v3.2.6), the maximum likelihood method (**B**.) implemented in the PhyML program (v3.1/3.0 aLRT) with 100 bootstrap replicates, and the neighbor joining method (**C**.) implemented in the BioNJ program. All trees were reconstructed with default settings and the graphical representation and edition of the phylogenetic tree were performed with TreeDyn (v198.3). Proteins and tree branches are depicted in blue for reptiles red for amphibians and in green for mammals. In panel **C**., the GenBank annotations are given for all protein sequences (with the exception of Shbg protein in *Anguilla anguilla* that were retrieved from the PhyloFish website), showing that all Shbga proteins are well annotated but that most Shbgb proteins are mis-annotated as Pros1-like or Gas6-like proteins.

**Supplementary Figure 2:**
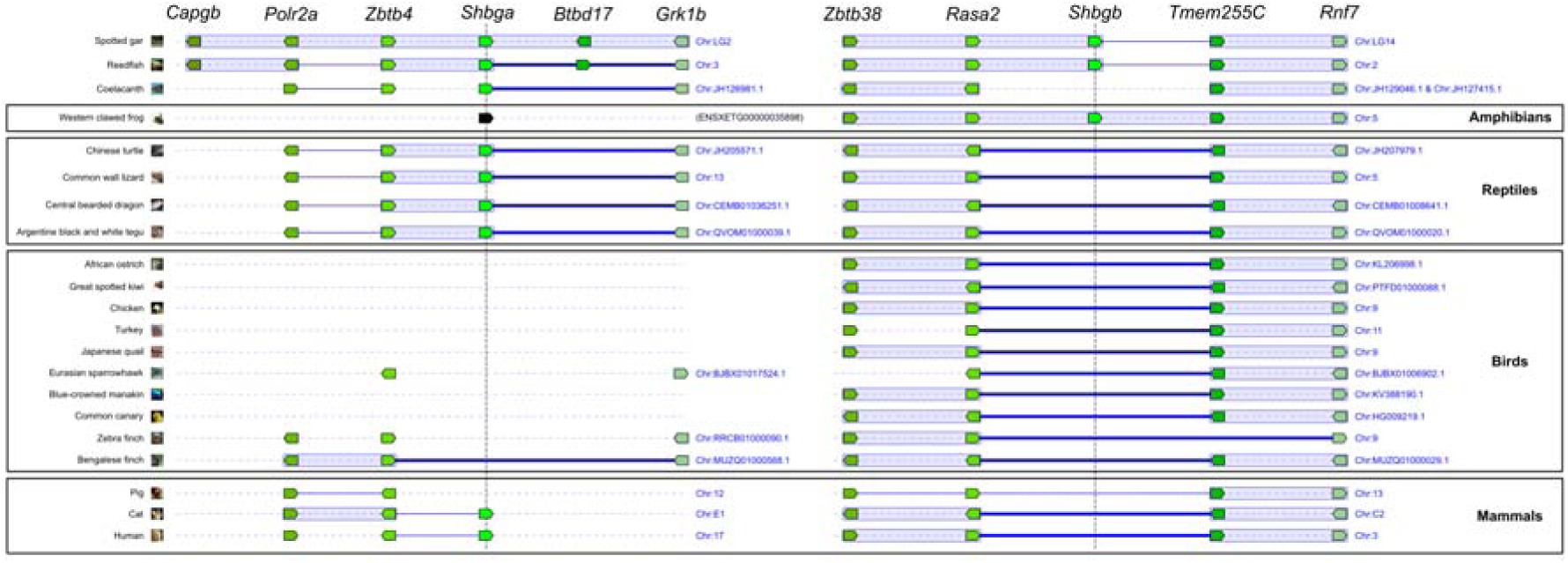
Genomic context and synteny relationships of some vertebrate Shbga and Shbgb proteins, showing the absence of both Shbga and Shbgb in birds and of Shbgb only in mammals and reptiles. This analysis was performed using the Genomicus genome browser (version 99.01, https://www.genomicus.biologie.ens.fr/genomicus-99.01/) using as seed sequences the accession numbers of Shbga (ENSLOCG00000013875) and Shbgb (ENSLOCG00000008107) of spotted gar (*Lepisosteus oculatus*). Only a subset of species is depicted in this figure but the absence of both Shbga and Shbgb in birds and of Shbgb only in mammals and reptiles was consistently found in all birds (N=35), mammals (N=99) and reptiles (N=14) genomes available in the Genomicus version 99.01. No Shbg homolog were found in the elephant shark, hagfish and lamprey genomes available in this Genomicus version 99.01.

